# Determinants of Severe Acute Malnutrition among Children age 6-59 Months Old in Two Public Hospitals, North West Ethiopia: a Case Control Study

**DOI:** 10.1101/664516

**Authors:** Adonia Damtew Nebro, Degnet Teferi Asres, Reddy PCJ Prasad

## Abstract

**Introduction:** Globally sever acute malnutrition affects 16.4 million under five children and more than one quarter of those children live in Africa. In Ethiopia, about 3% of children are severely wasted and continues to be persistent over the past 15 years. To implement an effective intervention, it is essential to identify predictors predispose to it. This study therefore, aimed to identify determinants of severe acute malnutrition among under five children in selected public health facilities, Northwest Ethiopia.

**Methods:** Institution based; age matched case control study was conducted on 104 cases and 208 controls. Bivariate and multivariate analyses were done using conditional logistic regression to identify predictors. Variables having P-value ≤ 0.2 during binary analysis were entered into multivariate analysis. P value < 0.05 was considered as statistically significant.

**Results:** Children from households of large family size(AOR=2.7, 95% CI: 1.06 – 6.9), having monthly income less than 1500 birr (AOR = 5.17, 95% CI: 1.7-15.3), which are food insecure (AOR = 2.9, 95% CI:1.17-7.28)), which didn’t receive any nutrition information (AOR= 3.47,95% CI: 1.14 - 7.10), didn’t practice exclusive breastfeeding (AOR = 2.69, 95% CI: 1.18 - 6.10), and practice infrequent hand washing (AOR= 7.6 95% CI:2.44-23.6) as well as children who had history of diarrhea two weeks prior to the survey (AOR 3.2, 95%CI:1.4-7.2) were more likely to suffer from severe acute malnutrition.

**Conclusion:** Family size, monthly income, food security status, exclusive breastfeeding practice, access to information on child feeding, hand washing practice and history of diarrhea were identified to be predictors of severe acute malnutrition. Due emphasis should be given to promoting family planning, improve household livelihoods and food security, strength awareness creation on exclusive breastfeeding and frequent hand washing practices as well as prevention of diarrhea.

## 1. Introduction

Severe acute malnutrition is defined as severe wasting (low weight for height) and/or mid-upper arm circumference (MUAC)<115 mm and/or bilateral pitting oedema (1).

Globally wasting is threaten the lives of 50.5 million under 5 children. Among this 16.4 million Children were severely wasted. Africa is the home of 13.8 million wasted children from which sever acute malnutrition threats the lives of 4 million children. At the same time the eastern Africa shares the burden of 1.5 million severely wasted children (2).

Children suffering from wasting an increased risk of death. Worldwide from 11.5% of the death occurred in children due to acute malnutrition 7.8% of the child death was attributed to sever acute malnutrition (3).

Despite the fact that the government of Ethiopia strives to increase its effort to enhance good nutritional practices through different programs, the poor nutritional status of children and women continues to be a serious problem. There has been a substantial decline in the proportion of children stunted and underweight. However, the proportion of wasted children remaining persistent overtime (4). According to the 2016 Ethiopian demographic & health survey report, 3 % of under five children are severely wasted. (5)

Similarly; Amhara region is among the highly affected region by malnutrition in Ethiopia. The proportions of children who are wasted and severely wasted were 9. 8% and 2.2% respectively (5). Higher prevalence of sever acute malnutrition (7.9%) were reported from a study done at Northwest Ethiopia(6). This indicates SAM is a serious public health problem in North West Ethiopia.

Study conducted in deferent part of the world shows that the predicators of acute malnutrition were inadequate care for children like poor infant and young child feeding, household food insecurity (7–9) incomplete/lack of immunization (10,11). poor environmental health condition (inadequate and unsafe water supply, and poor environmental sanitation) and mothers habit of less frequent hand washing and absence of latrine (7).

Though acute malnutrition is one of the public health problems and persistent over time in Ethiopia particularly in the study area, available studies on the determinants of SAM are limited. To implement an effective and efficient intervention, which can reduce the high burden of sever acute malnutrition, it is essential to understand factors predisposing to it in different part of the country. Therefore, this study is intended to assess the determinant factors of SAM in under five children in North West Ethiopia.

## Objective

- To identify predictors of severe acute malnutrition among children of 6-59 months of age in Felege Hiwot and Debre Tabor referral hospitals, Northwest Ethiopia.

## Methods

### Study design, period and setting

Institution based case control study design was employed from February 15, 2017 to June, 10 2017 at Felege Hiwot and Debre Tabor referal hospitals, Northwest Ethiopia. Felege Hiwot referral hospital is located in Bahir Dar. Bahir Dar (capital city of Amhara Regional State) and it is 565 km far to Northwest of Addis Ababa. It serves over 7 million people from the surrounding area. Debre Tabor referal hospital located in South Gondar zone of the Amhara Region of Ethiopia (Debre Tabor town) which is about 100 kilometers southeast of Gondar and 50 kilometers east of Lake Tana. It provides services for 2.3-2.5 million people of the surrounding area.

### Source and study population

All children 6–59 months of age with SAM or without malnutrition that have been admitted and treated at therapeutic feeding units (TFU) and/or other pediatric units of the two hospitals were the source population. The study population were 312 children age 6-59 months who were selected for this study from both hospitals during the study period.

### Inclusion and exclusion criteria

Children in the age of 6-59 months who were admitted in the two hospitals due to severe acute malnutrition with their care takers/mothers were included into the study as cases. Children aged 6-59 months with normal nutritional status and who had visited pediatrics units for different health care services during the study period were included as controls.

Children in the age of 6-59 months with disability (physical deformity and handicapped) and children who had sudden shock (unconscious) during the study period were excluded. Children with secondary undernutrition due to other pathological disorders and with other causes of edema were also excluded.

### Sample size and sampling procedure

The sample size was calculated using two population proportion formula, by taking maternal autonomy in decision making as major associated factor from other study and proportion of mother who had no autonomy in decision making to be 85.2% among cases and 69.4% among controls and detecting OR of 2.545 (12), with 95% CI, a power of 80%, a two to one ratio of control to case (2:1) and 10% non-response rate. By referring the above, the sample size was 276 (92case and 184 controls). But 104 cases and 208 controls were included assuming it could increase the power of the study.

The sample size was proportionally allocated to the two hospitals based on the average monthly follow rate of under five Children with severe acute malnutrition. Out of the total sample, 66 cases and 132 controls were allocated to Felege Hiwot referral hospital, while 38 cases and 76 controls were allocated to Debre Tabor referral hospital. Children in the age of 6-59 months admitted due to SAM in the therapeutic feeding units of the two hospitals during the study period were included until the calculated sample size was saturated. The controls were match to the case within ±2 months. Once a case child was admitted, his/her mother/caretaker was interviewed immediately and then the first two controls matched with one case of a similar age in months were randomly selected immediately after the admission of a case.

### Data collection and quality assurance procedures

The data were collected from all study participants using interviewer administered structured questionnaire. The questionnaires were originally prepared in English and then translated to local language, Amharic and translated back to English language to check consistency. The questionnaire comprised of different variables including socio-economic and demographic factors, child caring practices (feeding practice, immunization), household food security status and environmental health conditions.

Household food insecurity was assessed by using the nine standard Household Food Insecurity Access Scale (HFIAS) questionnaire developed by the Food and Nutrition Technical Assistance (FANTA) (13).

Dietary diversity scores (DDSs) were estimated via a 24-h recall method using the seven food groups. The food groups assessed were; Grains, roots or tubers; Vitamin A-rich fruit and vegetables; other fruits and vegetables; Flesh foods (Meat, poultry, fish and seafood); Eggs; Legumes, Pulses or nuts; milk and milk products (14).

In order to select the controls, weights of the children were measured in kg to the nearest 0.1 kg with minimum clothing and the height in (cm) to the nearest 0.1 cm. MUAC of the children were measured in cm to the nearest 0.1 mm and presence of bilateral pitting edema was checked by applying thumb pressure on both feet of the child. After all these above anthropometric measurements, if children were found to normal in nutritional status they were included as controls.

To assure the quality of the data, properly designed, pretested questionnaire was used. Training was given for data collectors and supervisor for two days.

### Variable Definitions

**Cases:** children with severe acute malnutrition whose MUAC≤11.5cm, WHZ < −3 SD or with bilateral pitting oedema.

**Controls:** children with normal nutritional (without edema or their MUAC >12.5 cm, their WHZ >-2 SD).

**Diet diversity: -** A child with a DDS of less than four was classified as having poor dietary diversity; or else, it was considered to have good dietary diversity.

### Household food security status ((13))

- Food secure - Household experiences none of the food insecurity conditions, or just experiences worry, but rarely
- Food insecurity - If household worries about not having enough food sometimes or often and or experiences other food insecurity conditions

### Data Processing and Analysis

Data were coded and entered to EPI info version 7.0 and transferred to Stata 14.0 for analysis. Frequencies and cross tabulation were calculated to describe the study population in relation to relevant variables. Conditional logistic regression was used to fit the data to identify the predictors for SAM. Both bivariate and multivariate logistic regression was computed to identify the determinants of severe acute malnutrition. Variables which had p-value of ≤ 0.2 in the bivariate analyses were entered in to multivariate analysis. P values < 0.05 were considered to declare the statistical significance.

### Ethical consideration

Ethical clearance was obtained from Review committee of Faculty of Chemical and Food Engineering, Bahir Dar Institute of Technology. The regional health bureau ethical review board approved and gave formal letter to both hospitals. Permissions to collect the data were obtained from medical directors of the two hospitals. Informed consent was obtained from the parents’/care givers of the children prior to taking necessary data.

## Results

### Socio-demographic characteristics of the Study subjects

A total of 312 (104 cases and 208 controls) participated with response rate of 100%. The mean age of the cases in months was 17.8(SD ± 11.51) and it was 17.62 (SD ± 10.88) among the controls. The mean age of the mothers of cases and controls in years were 31.9(SD ± 6.04) and 30.01(SD ± 5.45) respectively. Regarding the sex of the children almost half (52.95%) of the cases and 54.8% of controls were male. About 64.4% of households of the cases and 46.6% of controls had the family size of > 4. Families of the cases who had monthly income of <1500 birr were 69.2% while the families of controls who had monthly income <1500 birr were 21.1% (Table 1).

**Table 1:** Socio-demographic and economic characteristics of the study participants in Felege Hiwot and Debre Tabor referral hospitals, February-June 2017

| Variables |  | Case n (%) | Control n (%) |
| --- | --- | --- | --- |
| Religion | Orthodox | 89(88.6) | 186(89.4) |
|  | Muslim | 15(14.4) | 22(10.6) |
| Age of the respondent | 15-34 | 71(68.3) | 180(76) |
|  | 35-58 | 33(31.7) | 50(24) |
| Residence | Urban | 32(30.8) | 123(59.1) |
|  | Rural | 72(69.2) | 85(40.9) |
| Head of household | Male headed | 96(92.3) | 201(96.6) |
|  | Female headed | 8(7.7) | 7(3.4) |
| Marital status | Married | 96(92.3) | 202(97.1) |
|  | Other | 7(6.7) | 3(1.4) |
| Maternal education | No formal education | 77(74.0) | 97(46.6) |
|  | Formal education | 27 (26) | 111 (53.4) |
| Paternal education | No formal education | 67(69) | 77(37.5) |
|  | Formal education | 28(28.86) | 129(62.9) |
| Paternal occupation | Government employ | 7(7.2%) | 60(29.3) |
|  | Farmer | 66(68.0%) | 83(40.5) |
|  | Merchant | 16(16.5%) | 46(22.4) |
|  | daily laborer | 8(8.2%) | 16(7.8) |
| Maternal occupation | Employed | 6(5.8) | 41(19.7) |
|  | Housewife | 24(23.1) | 55(26.4) |
|  | Farmer | 60(57.7) | 77(37.0) |
|  | Merchant | 14(13.5) | 35(16.8) |
| Monthly income (Birr) | <1500 | 72(69.2) | 44(21.1) |
|  | ≥1500 | 32(31.8) | 164(78.9) |
| Decision making to use money | Mostly mother | 20(19.2) | 25(12) |
|  | Mostly father | 61(58.7) | 93(44.7) |
|  | Jointly | 23(22.1) | 90(43.3) |
| Family size | <5 | 37(35.6) | 111(53.4) |
|  | ≥5 | 67(64.4) | 97(46.6) |
| Birth order | <3 | 44(42.41) | 118(56.7) |
|  | >3 | 60(57.69) | 90(43.3) |
| Under five children | One | 80(76.9) | 156(75.0) |
|  | Two | 24(23.1) | 52(25.0) |
| Sex of the child | Male | 55(52.9) | 114(54.8) |
|  | Female | 49(47.1) | 94(45.2) |
| Child Mean age in months (±SD) |  | 17.81(SD±11.5) | 17.62(SD±10.8) |
| Food security status | Food insecure | 66(63.5) | 44(22.6) |
|  | Food secure | 38(36.5) | 161(77.4) |

### Maternal and child caring practices

High proportions of the cases (64.4%) were not exclusively breastfed for the first 6 months of age compared to the controls (35.5%). About 64.42% of the cases and 23.1% of the controls had diarrhea to two weeks prior to the survey. Majority (95.1%) of mothers /caretakers of the cases and 87.1% of mothers/caretakers of the controls used to take their children to health facilities after 24 hour of the onset of diseases. About 87.65% of mothers/caretakers of the cases and 77.03% of mothers/caretakers of the controls didn’t give additional food when the children were sick (Table 2).

**Table 2.** Maternal and child caring practice of respondents at Felege Hiwot and Debre Tabor Hospitals, February – June 2017, n=312

| Variables |  | Case No (%) | Control No (%) |
| --- | --- | --- | --- |
| Exclusive breast feeding | **Yes** | 37(35.5) | 129(62.2) |
|  | **No** | 67(64.5) | 79(37.8) |
| Dietary diversity score | **< 4 food groups** | 64(61.5) | 63(30.7) |
|  | **≥ 4 food groups** | 40(39.5) | 145(69.3) |
| Age appropriate immunization | **Yes** | 96(92.3) | 202(98.7) |
|  | **No** | 8(7.6) | 6(2.3) |
| Child vitamin-A supplementation in the past six months | **Yes** | 58(49.3) | 151(47.9) |
|  | **No** | 46(50.7) | 57(52.1) |
| History of diarrheal in the previous two weeks the survey | **Yes** | 67(64.42) | 48(23.07) |
|  | **No** | 37(35.58) | 160(76.9) |
| History of fever in the past two weeks | **Yes** | 54(51.9%) | 106(51%) |
|  | **No** | 50(48.1) | 102(49%) |
| Take to health institution when the child is sick | **Within 24 hr** | 4(4.9%) | 19(12.9%) |
|  | **After 24 hr** | 77(95.1%) | 129(87.1%) |
| Nutrition information on child feeding | **Yes** | 51(49) | 172(82.69) |
|  | **No** | 53(51) | 36(17.31) |
| Place of delivery | **Home** | 24 (23) | 25(12.1) |
|  | **Health institutions** | 80(77) | 183(87.9) |
| ANC | **Yes** | 90(88.23) | 197(97.6) |
|  | **No** | 12(11.77) | 11(2.4) |
| Give additional food to sick child | **Yes** | 10(12.35) | 34(12.97) |
|  | **No** | 71(87.65) | 114(77.03) |

### Environmental health conditions

The families who had access to protected water source were 54.8% in the cases and 76.4% in the controls. Almost all households (97.1%) of the cases and 97.65% of the controls didn’t treat water at home. Frequent hand washing practice (before serving/preparing food, after defecation before feeding the child) was practiced in 61.5% of the mothers/caretakers of the cases and 89.4% of controls. Mothers who used to wash their hands using water only were 54.8% in cases and 31.2% in controls (Table3).

**Table 3:** Environmental health conditions of the respondents in Felege Hiwot and Debre Tabor Hospitals, February-June, 2017 n=312

| Variables |  | Case No (%) | Control No (%) |
| --- | --- | --- | --- |
| Source of water | Protected | 57(54.8) | 159(76.4) |
|  | Unprotected | 47(50.7) | 49(23.7) |
| Water treatment at home | Yes | 3(2.9) | 5(2.4%) |
|  | No | 101(97.1) | 203(97.6) |
| Waste disposals | Appropriate | 38(36.5) | 154(74.03) |
|  | Inappropriate | 66(63.5) | 54(25.7) |
| Hand washing practice of mothers/ caregivers | Wash frequently | 64(61.5) | 106(89.4) |
|  | Wash less frequently /not wash | 40(38.5) | 22(10.6) |
| Use of detergents or local cleansing gents | water Only | 57(54.8) | 65(31.2) |
|  | Sometimes with soap | 41(39.4) | 95(45.7) |
|  | Always using soap | 6(5.8) | 48(23.1) |
| Toilet facility | Yes | 60(57.69) | 183(87.98) |
|  | No | 44(42.31) | 25(12.12) |

### Determinants of sever acute malnutrition

The socio-demographic and economic characteristics that were significant predictors of SAM in the multivariable analysis include monthly income, family size, and household food security status. Furthermore, exclusive breastfeeding practice, access to information on child feeding and history of diarrhea two weeks prior to the survey and hand washing practice were also significant determinants.

Children from households with large family size (>5) were 2.7 times more likely to be affected by SAM as compared to their counterparts (AOR=2.7, (95% CI: 1.06 – 6.9). Children from families who had monthly income of < 1500 birr were nearly 5 times more likely to develop severe acute malnutrition as compared to children from families with monthly income of ≥1500 (AOR = 5.17, 95% CI:1.7-15.3). Children from food insecure household were also about 3 times more likely develop SAM as compared to their counterparts (AOR= 2.9, 95% CI: 1.17-7.28).

The odd of SAM was higher among children who were not exclusively breastfed compared to exclusively breastfed children (AOR = 2.69, 95% CI: 1.18-6.10). Children from mothers/care givers who didn’t receive nutrition information were 3.4 times more likely to be severely malnourished than their counterparts (AOR=3.47, 95% CI: 1.14-7.10).

Children whose mothers/care takers didn’t practice frequent hand washing were 7.6 times more likely to be affected by SAM than those who did (AOR 7.6, 95% CI: 2.44-23.6). Similarly, children who had history of diarrhea two weeks prior to the survey were about 3 times more likely to develop SAM as compared to their counterparts (AOR 3.2,95%CI:1.4-7.2)(Table 6).

**Table 6.** Both bivariate and multivariable resulted of determinants of sever acute malnutrition in Felege Hiwot and Debre Tabor Hospitals, February-June, 2017 (n=312)

| Variable |  | Case No (%) | Control No (%) | Crude OR |  |  | Adjusted OR |  |  |
| --- | --- | --- | --- | --- | --- | --- | --- | --- | --- |
|  |  |  |  | OR | 95% CI | P –value | OR | 95% CI | P – value |
| Residence | Urban | 32(30.8) | 123(59.1) | 1 |  |  | 1 |  |  |
|  | Rural | 72(69.2) | 85(40.9) | 3.48 | 2.03–5.97 | 0.000 | 1.56 | .328-7.42 | 0.567 |
| Maternal education | No formal education | 77(74.0) | 97(46.6) | 3.47 | 1.98 -6.08 | .000 | .79 | .204-3.07 | 0.736 |
|  | Formal education | 27 (26) | 111 (53.4) | 1 |  |  | 1 |  |  |
| Paternal education | No formal education | 67(69.07) | 77(37.1) | 4.5 | 2.51-8.11 | .000 | 2.5 | .638 - 9.88 | 0.187 |
|  | Formal education | 28(28.86) | 129(62.9) | 1 |  |  | 1 |  |  |
| Paternal occupation | Government employ | 7(7.2%) | 60(29.3) | 1 |  |  | 1 |  |  |
|  | Farmer | 66(68.0%) | 83(40.5) | 6.73 | 2.72-16.65 | 0.000 | .89 | .107- 7.35 | 0.914 |
|  | Merchant | 16(16.5) | 46(22.4) | 2.57 | .92 - 7.17 | 0.072 | 3.8 | .729-20.32 | 1.56 |
|  | daily laborer | 8(8.2%) | 16(7.8) | 4.92 | 1.45 -16.74 | 0.011 | 2.8 | 0.31-25.66 | 0.93 |
| Maternal occupation | Employed | 6(5.8) | 41(19.7) | 1 |  |  | 1 |  |  |
|  | Housewife | 24(23.1) | 55(26.4) | 2.83 | 1.06 -7.56 | 0.038 | .44 | .072 - 2.74 | 0.384 |
|  | Farmer | 60(57.7) | 77(37.0) | 5.46 | 2.14 - 13.93 | 0.000 | .20 | .022 - 1.76 | 0.148 |
|  | Merchant | 14(13.5) | 35(16.8) | 2.56 | 0.88 -7.42 | 0.083 | .48 | 0.59 - 3.85 | 0.491 |
| Monthly income(Birr | <1500 | 72(69.2) | 44(21.1) | 11.89 | 5.62-25.13 | 0.000 | 5.17 | 1.74 -15.3 | 0.003 |
|  | >=1500 | 32(31.8) | 164(78.9) | 1 |  |  | 1 |  |  |
| Decision making to use money | Mostly mother | 20(19.2) | 25(12) | 4.31 | 1.83 10.17 | .001 | .610 | .165-2.25 | 0.459 |
|  | Mostly father | 61(58.7) | 93(44.7) | 2.79 | 1.54-5.04 | .001 | .472 | .108-1.69 | .226 |
|  | Jointly | 23(22.1) | 90(43.3) | 1 |  |  | 1 |  |  |
| Family size | <5 | 37(35.6) | 111(53.4) | 1 |  |  |  |  |  |
|  | ≥5 | 67(64.4) | 97(46.6) | 1.92 | 1.14-3.24 | 0.014 | 2.7 | 1.06-6.93 | 0.036 |
| Food security status | Food insecure | 66(63.5) | 44(22.6) | 7.24 | 3.85-13.61 | .000 | 2.9 | 1.17-7.28 | 0.021 |
|  | Food secure | 38(36.5) | 161(77.4) | 1 |  |  | 1 |  |  |
| Exclusive breast feeding | Yes | 37(35.5) | 129(62.2) | 1 |  |  | 1 |  |  |
|  | No | 67(64.5) | 79(37.8) | 3.5 | 2.04-6.11 | .000 | 2.6 | 1.18-6.10 | 0.018 |
| Dietary diversity score | < 4 food groups | 64(61.5) | 63(30.7) | 4.18 | 2.13-6.33 | 0.00 | 1.32 | 0.488-3.61 | 0.56 |
|  | ≥ 4 food groups | 40(39.5) | 145(69.3) | 1 |  |  | 1 |  |  |
| History of diarrheal in the previous two weeks | Yes | 67(64.42) | 48(21.58) | 4.57 | 2.70 - 7.35 | .000 | 3.2 | 1.40-7.28 | 0.006 |
|  | No | 37(35.58) | 159(78.42) |  |  |  | 1 |  |  |

| Continued... |  |  |  |  |  |  |  |  |  |
| --- | --- | --- | --- | --- | --- | --- | --- | --- | --- |
| Child vitamin-A supplementation | Yes | 58(49.3) | 151(47.9) | 1 |  |  | 1 |  |  |
|  | No | 46(50.7) | 57(52.1) | 2.50 | 1.44-4.47 | 0.001. | 1.19 | .44-3.26 | 0.72 |
| Place of delivery | Home | 24 (23) | 25(12.1) | 2.4 | .1.26-4.69 | 0.000 | .77 | 2.57-.230 | .641 |
|  | Health institutions | 80(77) | 183(87.9) | 1 |  |  | 1 |  |  |
| Nutrition information on child feeding | Yes | 51(49) | 172(82.69) | 1 |  |  | 1 |  |  |
|  | No | 53(51) | 36(17.31) | 5.7 | 3.22-10.13 | 0.000 | 3.4 | 1.5-7.9 | 0.003 |
| Source of water | Protected | 57(54.8) | 159(76.4) | 1 |  |  | 1 |  |  |
|  | Unprotected | 47(50.7) | 49(23.7) | 2.74 | 1.62-4.63 | 0.000 | .345 | .109-1.08 | .070 |
| Waste disposals | Burning /pit | 38(36.5) | 154(74.03) | 1 |  |  | 1 |  |  |
|  | Open field | 66(63.5) | 54(25.7) | 5.9 | 3.25-10.74 | 0.000 | 1.46 | .513-4.00 | 0.492 |
| Hand washing practice of mothers | Wash frequently | 64(61.5) | 106(89.4) | 1 |  |  | 1 |  |  |
|  | Wash less frequently/not wash | 40(38.5) | 22(10.6) | 5.5 | 2.89-10.74 | 0.000 | 7.6 | 2.44-23.6 | 0.000 |
| Use of detergents or local cleansing agents | Water only | 57(54.8) | 65(31.5) | 9.3 | 3.14-27.55 | 0.000 | 1.38 | 0.28- 4.49 | 0.658 |
|  | Sometimes with soap | 41(39.4) | 95(45.6) | 4.35 | 1.42-13.32 | 0.010 | 1.8 | .537-6.85 | 0.383 |
|  | Always using soap | 6(5.7) | 48(22.9) | 1 |  |  | 1 |  |  |
| Toilet facility |  |  |  |  |  |  |  |  |  |
| Yes |  | 60(57.69) | 183(87.98) | 1 |  |  | 1 |  |  |
| No |  | 44(42.31) | 25(12.12) | 5.55 | 2.96-10.42 | 0.000 | 2.71 | 1.06 – 6.93 | 2.10 |

## Discussion

Factors contributing to SAM are multifaceted and identifying the determinants of SAM in the study area can be very important to implement effective intervention. In this study, the determinants identified for SAM were family size, monthly income, household food security status, exclusive breastfeeding practice, hand washing practice and diarrhea history two weeks prior to the survey.

Accordingly, children from households with large family size (>5) were 2.7 times more likely to be affected by SAM as compared to children from household with smaller family size. The finding was in line with a study done in Sudan, which revealed that having more than four family members in the household was associated with acute malnutrition (15). This might be because of the fact that increased number of families placed a heavy burden on scarce household resources, particularly on financial and food. That might make difficult to fulfill the dietary need of the children. Besides, the increased number of family might reduce the time and quality of care received by the children.

Children from families who had monthly income of less than 1500 birr were nearly 5 times more likely to develop SAM as compared to children from families who had monthly income of 1500 birr and above. This finding was comparable with the study conducted in Jimma Zone, Ethiopia; which points out that children whose family monthly income of less than $50(1450 Ethiopian Birr) were 6 times more likely to be affected by SAM (16). This can be explained by the fact that households with low income could not afforded to buy food for consumption which results in inadequate diet that leads to child malnutrition.

The odd of severe acute malnutrition in children increased when the household was food insecure. Children from food insecure households were 3.7 times more likely to be affected by SAM than children whose families were food secured. This is consistent with a study done in west Gojjam, Northwest Ethiopia (17) and Haromaya District, Eastern Ethiopia (18). In contrary, a study from Gambela town, Western Ethiopia and Sekela District, Western Ethiopia revealed that there was no significant association between household food insecurity and acute malnutrition in children aged under 5 years(19,20). In this study, household food insecurity was found to be significant determinant of SAM and this might be due to limited availability of food or no economic access to purchase it which might lead to reduced quantity and quality of diet. Therefore, food insecure households might not satisfy the dietary need of household member, especially those under five children because they are at greater risk of malnutrition due to their higher nutrient demand for their growth. (21,22)

In this study exclusive breastfeeding practice was one of the determinants of SAM among children. Children who were not exclusively breastfed were nearly 4 times more likely to suffer from SAM. Similar findings were also documented in other studies done in Oromia Region and East rural Ethiopia (7,23). The study done in EAG States, India also indicated that children who were exclusively breastfed were found to be 16 percent less likely to be mild wasted and 48 percent less likely to be not wasted (24). This could be explained by the fact that children who were not exclusively breastfed had lower chance of preventing infections as breast milk has many immunological properties that are likely to protect against infections in infancy. A study done in Tanzania confirmed that exclusive breastfeeding reduce the risk of diarrhea dysentery(25). Besides, there might be contamination of bottles and foods that were early introduced to the child which contributes to higher risk of diarrheal disease in children.

Access to nutritional information on child feeding practice was also another determinant of SAM identified in this study. Children whose mothers/caretakers didn’t have any nutrition information on child feeding practice were nearly 3 times more likely to develop SAM than those children whose mothers/caretakers had information on child feeding practice. This finding is also supported by other study which is done in Botswana (26). Lack of information on child feeding might lead to inappropriate child feeding practice which then could affect the nutritional status of the children.

Hand washing practice was another factor that was found to be a significant determinant of SAM. Children whose mothers didn’t wash their hands before serving food, after defecation, before feeding the child and after cleaning the child were 7 times more likely to develop SAM as compared to children whose mothers washed their hands in each activity. Similar findings were documented in studies conducted in Oromia region, West Ethiopia and Machakel district, Northwest Ethiopia (7,27). This might be due to the fact that poor hand washing practice might lead to contamination of foods and that might increase a risk of infections and diarrheal diseases that in turn might lead to poor appetite and poor absorption of nutrient and finally might expose the child to SAM.

Moreover, history of the diarrhea two weeks prior to the survey was identified to be significant predictor for occurrence of SAM. The odd of SAM increased by three fold in children who had diarrheal disease two weeks prior to the survey than those who didn’t. Similar findings were reported from studies conducted in Shashogo, southern Ethiopia and Oromia, West Ethiopia (7,12).This could be due to the fact that diarrheal disease might lead to loss of appetite, loss of nutrients from the body and poor absorption of nutrients consumed.

### Limitations of the study

Since the questions were relied on the memory of the mothers/caretakers, this might introduce recall bias. To minimize this bias the recalling period was made shorter for some variables like history of diarrhea within two weeks prior to survey and about food consumption during last 24 hours which is a reasonable recalling period. However, for some of the variables like exclusive breastfeeding practice and vaccination received might introduce recall bias. There might be also selection bias because controls were selected from health facilities.

## Conclusion

The findings of this study confirmed that family size, monthly income, food security status, exclusive breastfeeding practice, access to information on child feeding, history of diarrhea two weeks prior to the survey and hand washing practice were significant determinants of severe acute malnutrition. due attention should be given to improve the knowledge and practice of parents towards exclusive breastfeeding and family planning by integrate breastfeeding promotion and support throughout the maternal and child health continuum, particularly in the prenatal and postpartum periods. It is also important to strengthen prevention and control of diarrheal disease through promotion of good hygiene (hand washing) and exclusive breastfeeding practices in the community. Creating income generating activities to improve the income and food security status of the poor households is also highly recommended. Since determinants of SAM were identified to be multifaceted and it cannot be addressed by a single sector only, the government should build strong multi-sectorial collaboration (Health, WASH and Agricultural sectors) to address the problem of severe acute malnutrition.

## Acknowledgment

The authors acknowledge Bahir Dar Technology Institute, Bahir Dar University for funding the research. We would like to thank data collectors and women for their willingness to participate. We also thank Amhara regional health bureau, Felege Hiwot and Debre Tabor Referral hospitals for their support and cooperation.

